# Optimised haemoglobin depletion improves clinical proteomics from dried blood spots

**DOI:** 10.64898/2026.06.13.731967

**Authors:** Hannah Ging, Rosemary E. Maher, Elin Davies, Philip J. Brownridge, Anirudh Rao, Alan D. Salama, Louise Oni, Claire E. Eyers, Andrew J. Chetwynd

## Abstract

Equitable access to large sample cohorts for robust, high-throughput proteomics for biomarker discovery is a major barrier to widescale clinical implementation. Dried blood spots (DBS) offer a minimally invasive alternative to venous blood draws, enabling at-home microsampling (<50 µL) for centralised analysis, thus enhancing research participation. This approach is particularly relevant for under-represented groups, including children, the elderly, minority backgrounds and those with long-term health conditions such as chronic kidney disease (CKD), where disease fluctuations may occur outside the clinic, and vein preservation is critical. Proteomic analysis has demonstrated great utility in monitoring disease progression, and for biomarker/therapeutic target discovery. However, liquid chromatography–tandem mass spectrometry (LC–MS/MS) of whole blood is hindered by the wide dynamic range and the relatively high abundance of proteins such as haemoglobin, compromising biomarker discovery. Here, we establish an optimised workflow for protein extraction and haemoglobin depletion from microsamples obtained using DBS, enabling sensitive and high-throughput proteomic analysis. We demonstrate that haemoglobin depletion increases protein identifications by ∼50%, mitigating ion suppression and dynamic range effects, enabling the identification of putative biomarkers from patients with stage 5 CKD on dialysis. We also evaluated a commercial cell-free DBS device which yielded a sample more representative of plasma compared to traditional DBS and enabled greater depletion of haemoglobin compared to traditional DBS with haemoglobin depletion methods. Our findings offer a scalable approach for biomarker discovery, facilitating remote, longitudinal clinical studies.

## Introduction

The rapid advances seen in the precise genomic characterisation of genetic diseases has highlighted the need to advance the field of clinical proteomics to improve characterization of diseases that may be driven by, or caused by, protein-based abnormalities that do not necessarily have a genetic component (1, 2). Clinical proteomics thus shows significant promise for both advancing the understanding of disease pathogenesis and potentially highlighting novel therapeutic targets (3, 4, 5, 6, 7). Typically, clinical proteomics utilises fresh plasma or serum samples that are processed on site and stored for batch analysis, requiring specialised laboratory equipment and on-site phlebotomists, with potential for substantial variability depending on the local processing procedures (3, 8, 9).

There is growing interest in the use of ‘micro-sampling’, where the volume collected is typically 50 µL or less (10), to overcome these limitations. An example of micro-sampling is the collection of dried blood spots (DBS), typically from a capillary finger-prick. DBS offer several advantages over traditional venipuncture given their compatibility with remote, microvolume, longitudinal sampling, low cost, and the fact that they are significantly less invasive than venipuncture (11, 12). Studies also show that DBS are preferred to venipuncture by patient populations as they reduce the likelihood of negative reactions such as dizziness and sampling fatigue (13, 14). Additionally, such microsampling aligns with the “Save the Vein” initiative that is particularly important when monitoring long term health conditions such as chronic kidney disease (CKD) where preservation of vein health is important for the formation of arterio-venous fistula for haemodialysis access (15). DBS have been used to understand viral infection and drug dosing, following interrogation of the proteome and/or the metabolome (16, 17, 18, 19). However, wide-spread clinical application of DBS sampling is not yet routine, with a major challenge being the variability in interference from cellular components when blood cells haemolyse upon drying (20, 21), along with a lack of standardisation in sample processing methodologies such as DBS device and sample reconstitution (11).

Currently the two DBS devices most commonly in clinical use are the Whatman903 and Ahlstrom226 filter papers. However, these are not volumetric and therefore pose a risk of quantitative inaccuracy (22, 23). Volumetric sampling devices are now growing in popularity, allowing for more regulated collection of DBS, and eliminating the effects of varying haematocrit concentrations, with demonstrable application across a number of fields, particularly pharmacology (24, 25, 26, 27, 28), and ‘omics (29, 30, 31, 32, 33, 34).

A notable challenge in clinical proteomics is the broad dynamic range of protein concentration in blood products, with haemoglobin accounting for 95% of the protein load in whole blood (35), whilst albumin, globulins and fibrinogens are together reported to make up 99% of the protein concentration in plasma (36, 37). These high abundance proteins can compromise the ability to identify candidate biomarkers due to ion suppression of lower abundance analytes during electrospray (36, 38, 39). To overcome this barrier in standard serum/plasma clinical proteomics workflows, depletion of haemoglobin and other highly abundant proteins, such as albumin, has traditionally been achieved using affinity binding mechanisms (40, 41, 42). Depletion of haemoglobin from reconstituted DBS is thus perceived to be a crucial step in establishing a sensitive DBS proteomics workflow for clinical application, which has to date hampered their use as convenient biomarker discovery vehicles (16, 43, 44). An alternative approach is the use of “cell-free” DBS devices, from companies such as Biodesix (45), Cobas (46), and more recently Capitainer, all of which remove blood cells and thus haemoglobin. These devices primarily operate via microfluidic separation (47, 48, 49) although passive filtration is also used (50, 51). Their ability to remove haemoglobin-containing cells at the point of sampling could provide broader use of DBS for clinical investigation.

Here we have evaluated and optimised four different commercial DBS devices for sensitive, robust label-free quantitative proteomics, considering reconstitution buffer, micro-sampling device, differences between plasma and blood, and the impact of haemoglobin depletion, comparing with more routine, liquid based sample processing (with and without abundant protein depletion). As proof of principle, we applied our optimised workflow to explore the changes in the proteome profiles of DBS from a cohort of patients with stage five CKD (CKD5).

## Materials and Methods

Unless otherwise specified, all reagents were sourced from ThermoFisher Scientific, Waltham, MA, United States.

### Ethical and Regulatory Approvals

For method development, samples were collected from healthy participants at the University of Liverpool (IRAS project identification: 327208). A total of 10 patients with stage 5 CKD and 10 healthy control adults were recruited in the proof-of-principle clinical validation aspect of this project using the ethically approved GlomOmics study (REC 23/PR/0490).

### Participants and Biosample Collection

Healthy participants were consented and recruited to support the initial method development using fresh heparinised whole blood taken from two healthy adult controls - one male and one female, ages 34 and 38. Participants with CKD5 managed on hemodialysis at the Royal Liverpool University Hospital, (Liverpool, UK) were recruited for clinical proof-of-principle immediately prior to starting a hemodialysis session. Demographic data were collected on all participants and all patients provided informed written consent, with samples handled under the principles of the Declaration of Helsinki. In all steps of the method development, 5 replicates were analysed.

### Dried Blood-Spot Devices and Sample Collection

The DBS devices used in this study were: 10 µL volumetric B10 and Sep10 (Capitainer Ab, Solna, Sweden), Whatman903 (Sigma Aldrich, St Louis, MO, United States), and 10 µL VAMS Mitra devices (Neoteryx, Torrance, CA, United States). For healthy control DBS, venous blood was collected in lithium heparin vacutainers and 10 µL was pipetted directly onto the B10, and Whatman903 devices. Sampling for the 10 µL VAMS Mitra devices was achieved by holding the device to the surface of the blood sample until saturated. For Sep10 devices, 70 µL of blood was pipetted directly onto the device, which was then tilted to 90° for 2 seconds. After in-device cell filtration, 10 µL of plasma-like solution was deposited on a collection disk. Samples were then allowed to dry for three hours in a fume hood, or overnight for the Sep10 devices as per manufacturer’s instruction.

For all optimisation, freshly drawn blood samples were used for each experiment, accounting for inter-experiment variation, as each experiment used different blood samples. Proof of principle clinical samples were obtained using capillary sampling using a finger prick device (BD Microtainer™, Fisher Scientific, Loughborough, UK)

### Sample Reconstitution

DBS samples were reconstituted using four different solutions: i) 200 μL of 50 mM Ammonium Bicarbonate (AmBic) (16), ii) 10 mM Phosphate Buffered Saline (PBS) (52), iii) 0.1% (v/v) Trifluoroacetic Acid (TFA) (53) in HPLC grade water, or iv) 67% 100 mM AmBic and 33% 2,2,2-Trifluoroethanol (TFE) (29) and incubated for two hours in a ThermoMixer (ThermoFisher Scientific, Waltham, MA, United States) at 800 rpm at room temperature (RT). The reconstituted sample was centrifuged at 700 *g*, 4°C for 15 minutes. The supernatant was then removed and stored in a fresh 1.5 mL tube. Alternatively, 133 μL of 50 mM AmBic and 67 μL TFE was added to the DBS in a microfuge tube and sonicated (15 minutes, RT), before incubation for 30 minutes in a shaking ThermoMixer (1000 rpm, RT). Samples were centrifuged at 700 *g*, 4 °C for 15 minutes, the supernatant collected and stored in a fresh low bind 1.5 mL tube (29).

A lysis bead reconstitution was also evaluated whereby HPLC-grade water (500 μL) was added to the DBS and homogenised with CKMix beads (Bertin Technologies SAS, Montigny-le-Bretonneux, France) in a Minilys Personal Homogeniser (Bertin Technologies SAS, Montigny-le-Bretonneux, France) for 5 minutes at RT (54). To the lysed sample, 500 μL Tris HCl (pH 8.5) and 0.5% (w/v) sodium dodecyl sulphate (SDS) were added. The sample was sonicated (100 W Cavitek Sonicator, Cavitek, Allendale Ultrasonics, Hertfordshire, UK) on full power for 5 minutes at RT. The sample was centrifuged at 700 *g*, 4 °C for 15 minutes, the supernatant collected and stored in a fresh low bind 1.5 mL tube.

In all cases, reconstitution was performed directly prior to further sample processing and analysis.

### High Abundance Protein Depletion

To deplete haemoglobin, HemogloBind Depletion and NuGel-HemogloBind Depletion (both 20 µL reconstituted blood) were performed as per manufacturer’s instructions (see supplementary information).

HemoVoid depletion was performed as per manufacturer’s directions, with all measurements halved to account for the reduced blood volume of 150 µL of reconstituted sample (see supplementary information).

### Protein Digestion

Reconstituted samples (25 µL) were digested using a modified SP3 protocol, and samples were washed with ethanol (55, 56). Samples were reduced and alkylated at 95 °C for 5 minutes with Tris(2-carboxyethyl)phosphine (TCEP)/chloracetamide (CAA) (120 mM/480 mM) (ThermoFisher Scientific, Waltham, MA, United States), then digested on E3 and E7 Sera-Mag SP3 beads (Cytiva, Marlborough, MA, United States) using LysC (Promega, Madison, WI, United States) for 2 hours, followed by Trypsin (Promega, Madison, WI, United States) for 16 hours at a 50:1 Protein: Protease ratio for each protease, at 1400 rpm and 37 °C (see supplementary information). Following digest, samples were acidified with formic acid to a final concentration of 1% (v/v) and centrifuged for 5 minutes at 2300 *g*. The supernatant was taken and dried to dryness using a SpeedVac (Centrifuge: UNIVAPO – 150 ECH, Cooling unit: UNICRYO MC2L -60 °C, Vacuum pump: UNIVAC DQ4) prior to freezing at -80 °C until LC-MS/MS analysis. In all cases, 5 experimental replicates were processed.

### LC-MS/MS

The dried peptides were resuspended in 100 µL 0.1% (v/v) FA, before diluting to a final peptide concentration of 5 ng/µL, determined using a NanoDrop 2000c (ThermoFisher Scientific, Waltham, MA, United States). EvoTips (EvoSep, Odense, Denmark) were prepared as per the manufacturer’s instructions (see supplementary information).

Peptides were separated using an EvoSep One LC system, applying a 30 samples per day method (see supplementary methods) on a C18 EASY-Spray HPLC column (2 µm particle size, 100 Å pore size, 150 µm x 150 mm). Solvent A was water with 0.1% (v/v) FA and Solvent B was acetonitrile with 0.1% (v/v) FA, freshly prepared for each experiment. Separated peptides were analysed using a QEx HF Orbitrap in positive electrospray ionization (ESI) mode. The MS1 scan range of 350-2000 *m/z*, with an automatic gain control (AGC) of 3E6, a maximum fill time of 100 ms, and a mass resolution of 60,000 FWHM at *m/z* 200. MS2 was acquired using a top 18 data-dependent acquisition (DDA) approach, between 200 to 2000 *m/z*, with a normalised collision energy (NCE) of 29% at a resolution of 30,000, an AGC of 1E5, and a maximum injection time of 45 ms.

### Data Analysis

Data was processed using Proteome Discoverer v2.4.0.305 (ThermoFisher Scientific, Waltham, MA, United States), searching against the UniProt human proteome database [accessed November 2025], using the MASCOT search engine. Precursor ion error tolerance was set at 10 ppm, and fragment ion error tolerance at 0.01 Da. Fixed modifications were set for carbamidomethylation of cysteine, and variable modifications for oxidation of methionine. The false discovery rate (FDR) was set at 1% at the level of the peptide spectral match. Data was then processed, exclusion of proteins absent in ≥25% of samples across all conditions, median normalisation and knn=3 imputation, and visualised using RStudio version 4.2.0, to produce Scatter graphs (Dunn Test, Holm adjustment), Principal Component Analysis (PCA) scores plots, Upset Plots, and Volcano plots (utilising Limma, Q-value<0.05). Gene Ontology (GO) Analysis was performed using ClusterProfiler and visualised in RStudio v4.2.0.

## Results and Discussion

### Optimisation of Sample Reconstitution Buffer

To compare the depth of the DBS proteome across the three DBS devices five reconstitution methods were evaluated, employing either AmBic, PBS, Lysis Beads, TFA, or TFE (Figure 1). Proteome coverage was evaluated by considering the number and reproducibility of proteins identified, as well as the consistency of quantification across all replicates.

**Figure 1.**
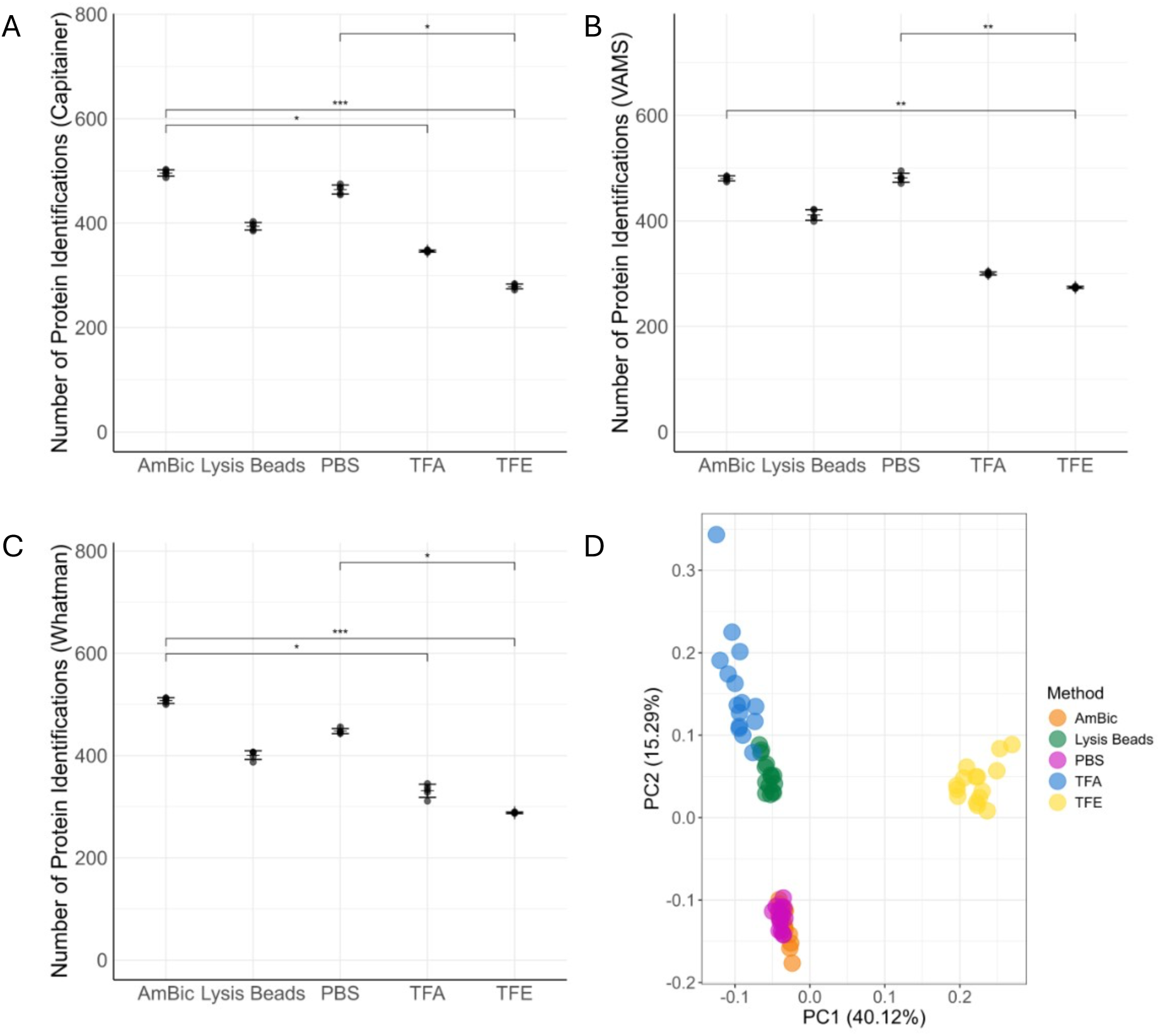
Comparison of reconstitution methods and DBS devices. (A) The average number of proteins identified in Capitainer b10 DBS with each reconstitution method (n= 5). (B) The average number of proteins identified in the Neoteryx VAMS Mitra DBS based on the reconstitution method (n= 5). (C) The average number of proteins identified in Whatman903 DBS with each reconstitution method (n= 5). (D) The data set (n= 5, total samples= 75) was subjected to PCA using the dried blood spot proteome, and colour coded by reconstitution buffer. *= p<0.05, **= p<0.01, ***= p<0.001, Error Bars= Standard Deviation. AmBic= Ammonium Bicarbonate, PBS= Phosphate Buffered Saline, TFA= Trifluoroacetic Acid, TFE= Tetrafluoroethylene.

No significant difference was observed in the total numbers of proteins identified using either AmBic or PBS as reconstitution buffers, and across all devices 495±13 and 465±16 proteins respectively were identified (Figure 1A, 1B, 1C). Slightly lower numbers of proteins were identified with the other reconstitution methods across all devices (Figure 1A, 1B, 1C) at 402±11, 326±21 and 280±7 for the lysis beads, TFA and TFE respectively. TFA yielded significantly fewer protein identifications than AmBic when using the Capitainer and Whatman devices (both p= 1.1x10^-2^). However, only the TFE reconstitution returned a statistically significantly lower number of protein identifications compared to both AmBic and PBS across all three tested devices. The number of proteins identified was reproducible for AmBic and PBS, with relative standard deviations (RSD) of 1.3% and 1.8%, respectively across all devices (Figure 1A, 1B, 1C). Across all three devices, protein identities were highly reproducible using both AmBic and PBS (Supplementary Figures 1 to 3) with over 90% of proteins being seen in all 5 replicates, for all three devices.

Reproducibility of protein quantification was assessed across all replicates per condition (Supplementary Figure 4). AmBic and PBS returned the lowest median RSDs of 1.23±1.28% and 1.34±1.72% respectively. This consistency was further highlighted through PCA visualisation, with replicates for AmBic and PBS extraction being tightly clustered (based on both PC1 and PC2) indicating the similarity of their identified proteomes (Figure 1D). Due to the numbers of proteins identified, the reproducibility of quantification, and compatibility with downstream processing pipelines, 50 mM AmBic was selected as the optimal reconstitution buffer.

### Comparison of Micro-Sampling Devices

In total, three DBS devices were assessed: b10, VAMS Mitra, and Whatman903. Although the highest number of proteins were identified using the Whatman903 device, at 508±6, compared to 496±6 and 481±5 for b10 and VAMS, respectively, all devices had RSD <1.5% showing remarkable reproducibility, and there were no significant differences in the number of proteins identified between the devices when replicated (n= 5) (p>0.999 for all) (Figures 1A, 1B, and 1C). Unlike the Whatman903 devices, VAMS and b10 devices are volumetric, accurately collecting a defined volume of blood, which supports reliability across the sample cohorts (22, 23). Furthermore, for shipping remote samples, the b10 and the VAMs devices provide a self-contained sample for convenience which is protected from damage and contamination.

Previous studies comparing the clinical performance of VAMS and b10 devices concluded that the b10 devices were the most consistent in terms of sampling success, with 92-96% of b10 samples being usable to monitor rejection drug concentration in transplant recipients (57). The study reported that comparatively usability of VAMS devices peaked at 72-88% when the sample was taken by a healthcare professional, and dropped 52-72% when the patient performed the sampling using the VAMs Mitra devices (57). Importantly, a patient engagement and acceptability study identified that the environmental impact of the micro-sampling device is an important consideration, and thus the primarily cardboard b10 devices provide a more environmentally sustainable option (14). As such, the b10 devices were selected for further optimisation.

As a negative control, blank DBS devices were also analysed to evaluate potential contamination. Following exclusion of those proteins that were present in <25% of samples including all devices, only 44 proteins were identified consistently across the three devices. Keratins made up 13 of these 44 proteins, accounting for ≈85% of the relative protein abundance, calculated using the intensities of each signal, with other proteins including albumin, dermcidin, actin, and two immunoglobulin proteins (For full list of identified proteins, see Supplementary Sheet 2).

### Comparison of Plasma, Dried Plasma and Dried Blood

To enable routine clinical sampling for proteomics, it is important to determine how closely the protein composition of DBSs represent the current gold standard of plasma. Comparing dried plasma spots with DBS (both processed in the same manner) and traditional plasma samples, we identified 392 proteins unique to the DBS (Wet plasma= 181±3, Dry plasma= 209±3, Dry blood= 573±2, p= 1.2x10^-3^) (Figure 2A and B), primarily due to the presence of cellular proteins in whole blood DBS (58). Gene ontology (GO) analysis confirmed that most of the proteins unique to DBS are cellular compartment proteins, particularly cytosolic proteins (Figure 2C) (Supplementary Sheet 3). These data mirror the findings of Molloy et al, 2022, who identified similar differences in protein number and composition between dried plasma and blood samples using VAMS devices (43).

**Figure 2.**
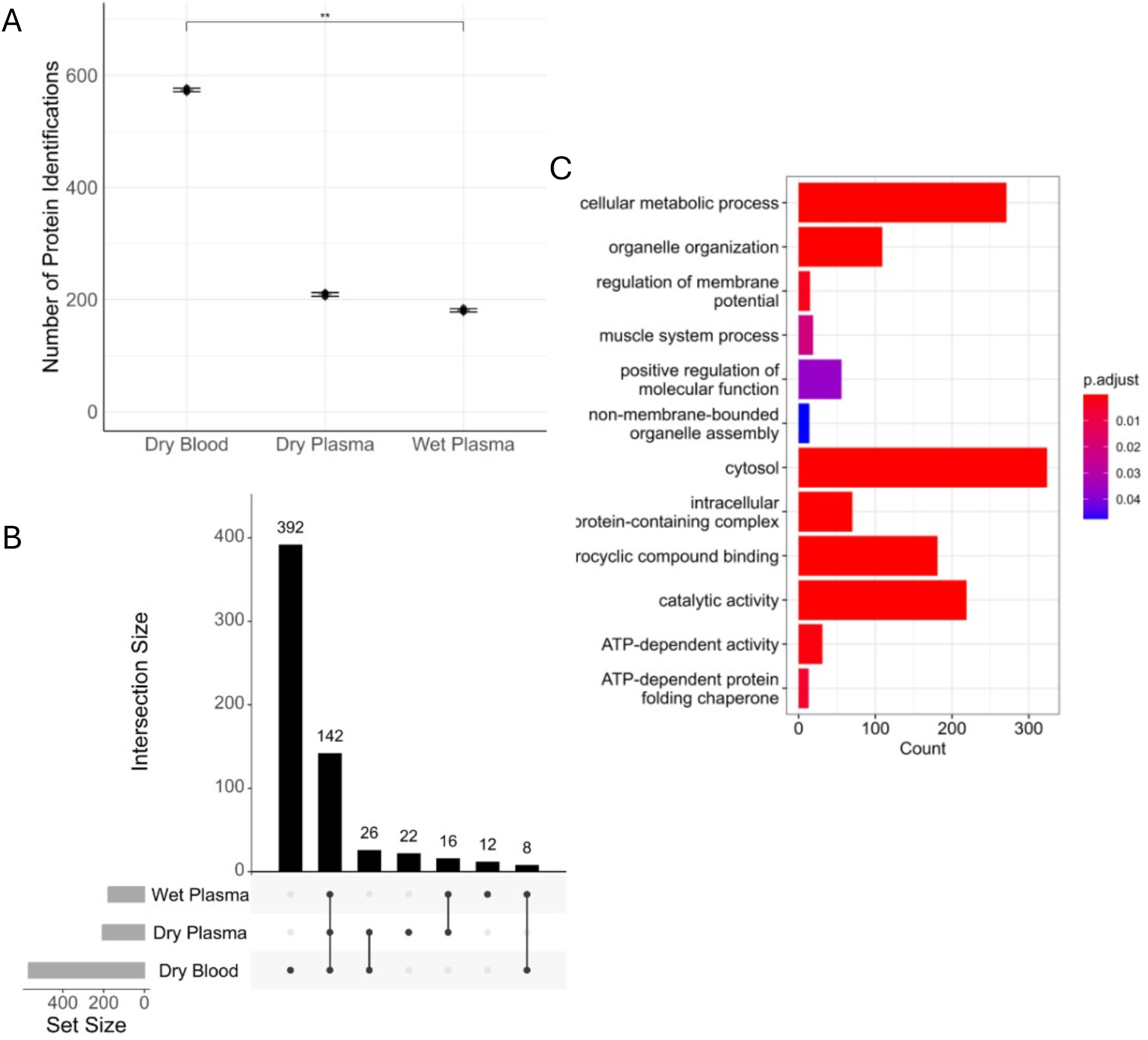
Comparison of DBS with dried plasma and standard plasma collection (A) The average number of proteins identified in dried blood spots, dried plasma spots, and wet plasma. (n= 5). (B) Upset plot exploring the similarities and differences between the proteins identified across sample types. (C) Gene ontology analysis of proteins only found to be present in the dried blood spot proteome, when compared to both the dried plasma spot and wet plasma proteomes. **= p<0.01, Error Bars= Standard Deviation.

There was <15% difference in the number of proteins identified using wet versus dry plasma, showing no statistical significance (p= 0.15) (Figure 2A, Supplementary Sheet 4). Twelve proteins were exclusively identified in the wet plasma samples, of which five were immunoglobulins or apolipoproteins which would typically be removed via high-abundance protein depletion methods. Of the remaining seven proteins, two were of particular clinical interest: peptidase inhibitor 16 (PI16), and cystatin-C (CST3). PI16 and CST3 have both been implicated as markers for different disease states; loss of CST3 is used as a measure of kidney function in patients with CKD, although its expression can also be altered by other factors such as obesity (59), and PI16 has been implicated in diseases such as diabetes (60).

A study by Chambers et al, 2013, also explored the effects of the ‘drying phase’ on blood-derived biofluids, using Whatman903 filter papers for dried serum, plasma and blood samples, compared to ‘wet’ controls. Chambers’ study did not identify any significant differences in the number of proteins identified between the dried and control conditions for any of the three biofluids, supporting the findings presented here (61).

### Evaluation of High Abundance Protein Depletion Techniques

Historically, exploration of the whole blood proteome has been hindered due to haemoglobin suppressing the signal from proteins of lower abundance (35). This has resulted in the identification of only limited portions of the whole blood proteome, with DDA-MS of DBS identifying protein numbers ranging from 120 to 350, and the high abundance of haemoglobin being cited as the leading cause for low proteome coverage (16, 43, 44). Therefore, methods to remove haemoglobin from DBS are key to the adoption of DBS sampling. We therefore evaluated three affinity-based methods (NuGel, HemogloBind, and HemoVoid) to deplete haemoglobin for improved DBS proteome coverage. Processing time for all three bead-based methods was similar, following very similar processes. However, while NuGel and HemogloBind deplete haemoglobin directly by affinity binding, capturing the depleted proteome, the HemoVoid beads have very low affinity for haemoglobin, which instead act to capture and enrich lower abundant proteins. As shown in Figure 3A, this results in a statistically significant increase in the number of proteins identified using HemoVoid compared with the other sample preparation devices (or undepleted blood). The increasing numbers of proteins identified from the NuGel (382±10), HemogloBind (423±5), and HemoVoid (545±5) compared with undepleted sample (364±5), inversely correlates with the proportion of haemoglobin removed (Figure 3B), potentially due to a lower dynamic range and reduced suppression during peptide ionisation: NuGel was the least efficient for haemoglobin removal (63±20.7% remaining), followed by HemogloBind and HemoVoid, which respectively retained 40±28.4% and 24±7.9% of the haemoglobin (p= 5x10^-3^ compared to undepleted samples) (Figure 3B). As well as enabling a 50% increase in the number of proteins identified compared with undepleted material, reproducibility of identification was also greatest with HemoVoid, with 94.7% of proteins being identified in all five replicates (Supplementary Figure 5). It is also worth noting that only 82 out of the 624 proteins identified across all experiments were not identified using HemoVoid (Supplementary Figure 6 and Sheet 6).

**Figure 3.**
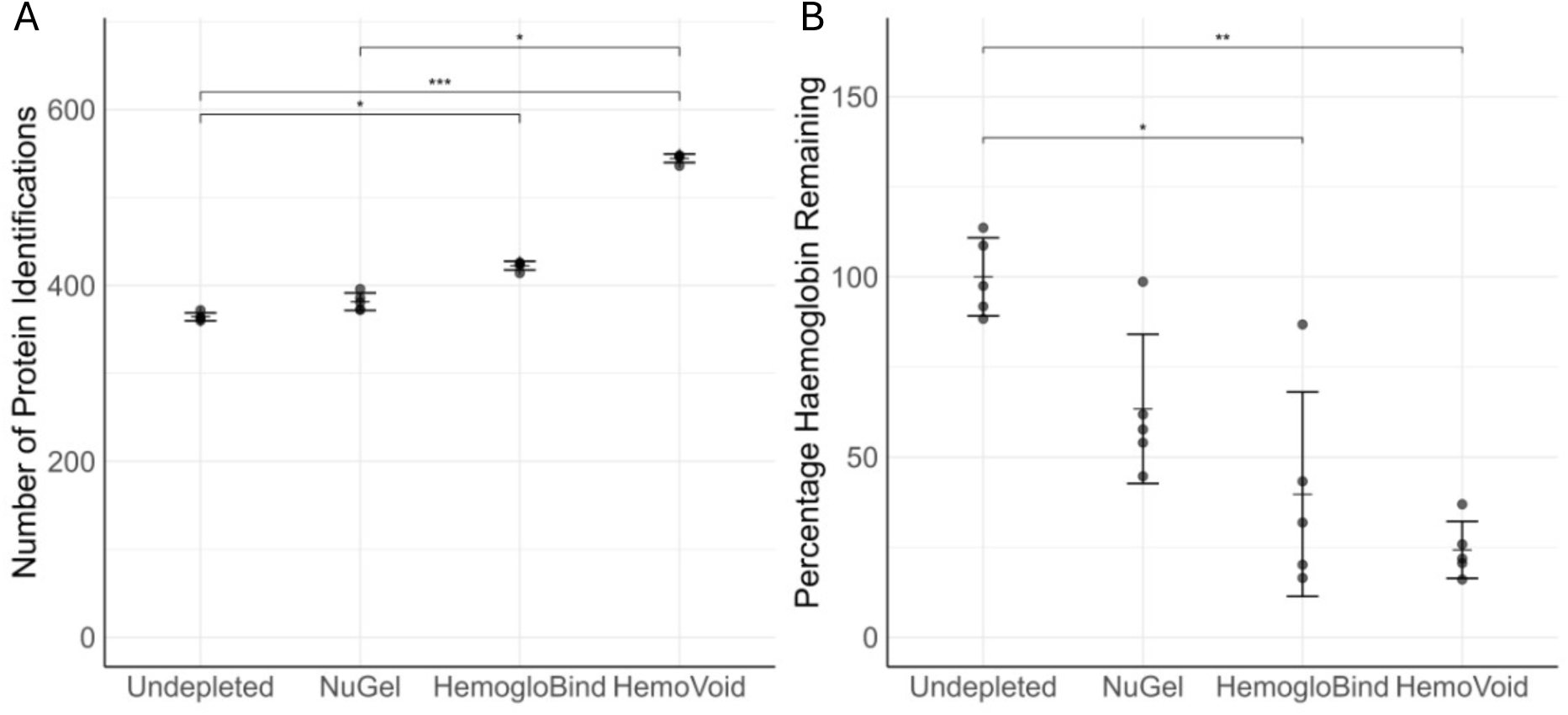
Comparison of haemoglobin and high abundance protein depletion methods. (A) The average number of proteins identified in the dried blood spot proteome for each depletion method (n= 5). (B) The average percentage of haemoglobin depleted from DBS calculated based on the undepleted condition (n= 5). *= p<0.05, **= p<0.01, ***= p<0.001, Error Bars= Standard Deviation.

### Cell Free Dried Blood Spot Analysis

An alternative method to collect haemoglobin-free DBS is a cell free collection device (47). These devices produce a “plasma like” sample by filtration, which is then dried *in-situ*. In this study, Sep10 devices were used, which filter out both the red and white blood cells. Removal of cellular components during filtration has the added advantage of also removing red blood cell-containing haemoglobin. Performance of the Sep10 devices was compared with conventional DBS collected using b10 devices, dried plasma spots (b10), and wet plasma samples. The DBS samples underwent HemoVoid depletion prior to analysis.

As expected, due to the presence of cellular material, significantly more proteins were identified in the DBS samples (386±2, RSD= 0.44%) compared with wet plasma (156±1, RSD= 0.45%, p= 1.1x10^-3^) (Figure 4A). The number of proteins identified using the Sep10 devices (164±4, RSD= 2.18%) was not significantly different from the wet plasma samples (p= 0.15), although the Sep10 data exhibited greater variability, albeit within acceptable limits (<5%). These results indicate that cell-free DBS collection offers a reliable approach for sample collection without the need for additional haemoglobin depletion (Supplementary Figure 7).

**Figure 4.**
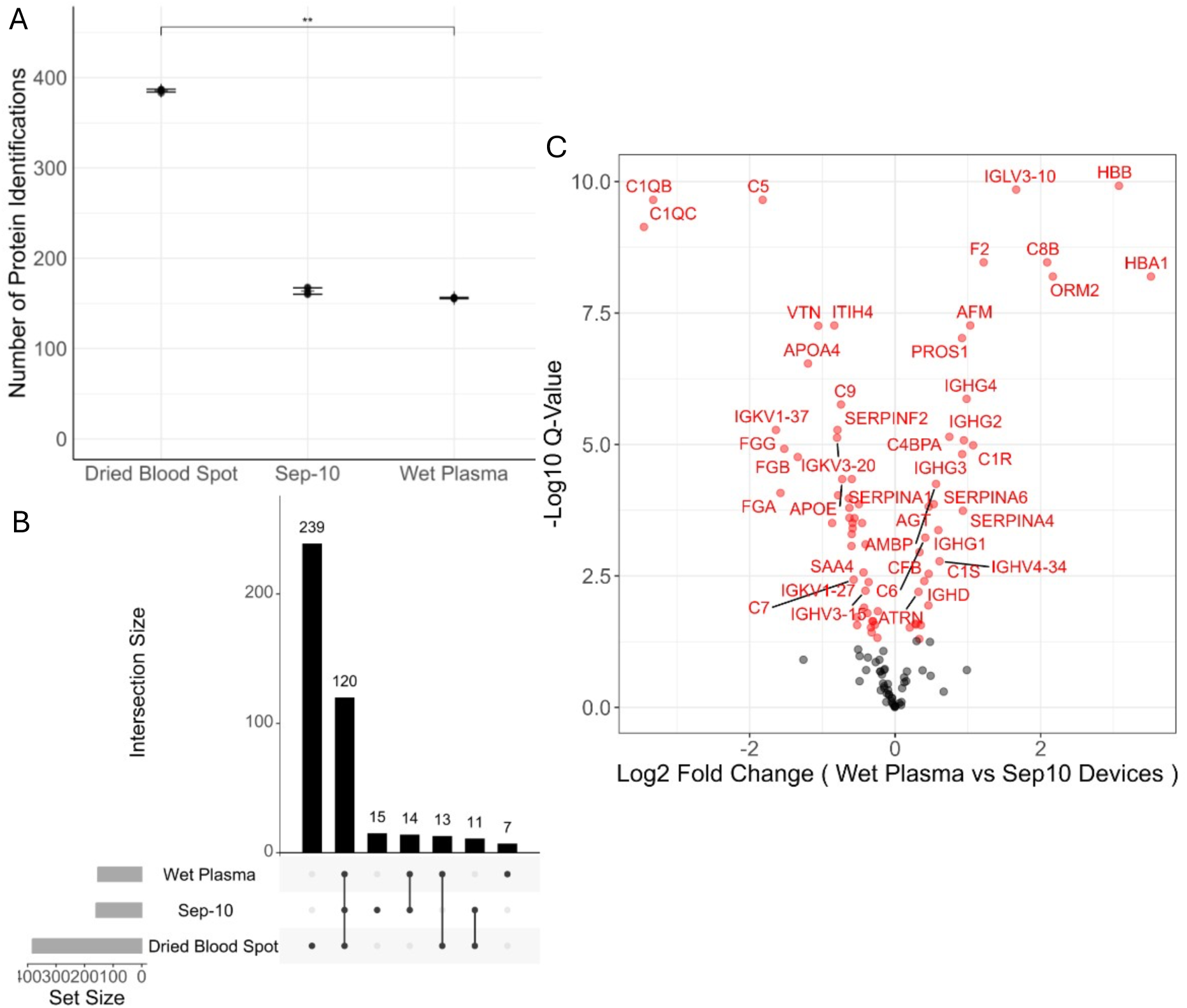
Analysis of cell-free DBS compared to traditional DBS and plasma samples. (A) The average number of proteins identified in each sample (n= 5). (B) Upset plot showing the cross over of proteins identified in each sample type (n= 5) (C) Volcano plot displaying log2mean fold change in protein abundance against -log10, Q-values derived from 2-Sample significance tests (limma) comparing the identified proteomes from wet plasma (left) to Sep10 devices (right). **= p<0.01, Error Bars= Standard Deviation. Protein names correlating to the Gene Name identifications can be found in the Supplementary information (supplementary table 1).

Of the 383 proteins consistently identified using the b10 devices with HemoVoid depletion in this experiment, 120 were not identified using the Sep10 devices (Figure 4A, 4B). This difference can be attributed to the acellular nature of the Sep10-derived matrix and resulting absence of cellular proteins from both the red and white blood cells, as previously discussed. A complete list of proteins unique to DBS samples is provided (Supplementary Sheet 7).

No significant difference was observed in the overall number of proteins identified using the Sep10 devices compared with plasma. However, there was a statistically significant difference in the relative abundance of 73 proteins (of the 122 proteins identified in 75% of all samples) between these two methods (Figure 4C, Supplementary Sheet 8). Of these, 43/73 (59%) were higher in the plasma samples, and 30/73 (41%) in the Sep10 devices, with specific complement proteins and immunoglobulins being differentially abundant between the two sample preparation approaches. Notably, haemoglobin subunits HBA1 and HBB were both at significantly higher relative abundances is samples prepared using the Sep10 devices than in plasma, suggesting reduced efficiency of haemoglobin and red blood cell removal than traditional centrifugation methods. This may be due to cell lysis during the Sep10 filtering process.

### Proof-of-Principle Clinical Validation in Chronic Kidney Disease

To demonstrate proof-of-principle for application of the developed HemoVoid method for proteome interrogation in a complex clinical disease state, we analysed samples from 10 patients with CKD5 undergoing haemodialysis alongside 10 healthy controls (*Table 1*), comparing with standard plasma. CKD patients were chosen due to the chronic nature of the disease, a growing economic and health burden, and the need to preserve vein health, highlighting the value of alternative approaches such as capillary sampling (15). For this clinical proof-of-principle study, HemoVoid was selected as the method of choice given the familiarity of whole blood DBS use in other biobanks, such as the Danish Newborn Screening biobank (62).

**Table 1.** Subject demographic and clinical characteristics. Dialysis vintage refers to the months since dialysis treatment commenced. All patients were diagnosed with Stage 5 chronic kidney disease (end stage kidney failure).

| Demographics |  |  |  |
| --- | --- | --- | --- |
|  | Chronic Kidney Disease (Stage 5) | Healthy control | Significance (p value) |
| Number of patients | 10 | 10 |  |
| Age (years, median and SD) | 73.5 ± 11.1 | 43 ± 7.9 | 4.5 x 10<sup>-6</sup> |
| Sex (% Male) | 50% | 60% |  |
| Dialysis vintage (months and SD) | 24.9±28.6 |  |  |
| Primary renal disease (n) | IgA Nephropathy 2<br>Granulomatosis with Polyangiitis 1<br>Diabetic Nephropathy 3<br>Renovascular disease 1<br>Obstructive Uropathy 1<br>Unknown 2 |  |  |
| Laboratory values |  |  |  |
| eGFR mL/min/1.73 m<sup>2</sup> (median and SD) | 5.5 ± 2 | 90 ± 6 | 4.2 x 10<sup>-19</sup> |
| C-reactive protein mg/L (median and SD) | 8.5 ± 7 | 1 ± 6 | 4.6 x 10<sup>-2</sup> |

Using this approach, we identified 702±16 proteins in the CKD5 cohort, compared with 685±13 in the healthy control DBS samples. Of these, 31 were present at significantly different levels between the two groups (Figure 5A, Supplementary Sheet 9), with 23 (74%) elevated in the CKD5 group and 8 (26%) higher in the healthy control cohort. This included CRP, which is a routinely used clinical marker of inflammation(63). CRP was significantly elevated in the CKD group (Q-value= 3.9x10^-2^, log2FC= 3.45), consistent with expectations for the chronic disease state.

**Figure 5.**
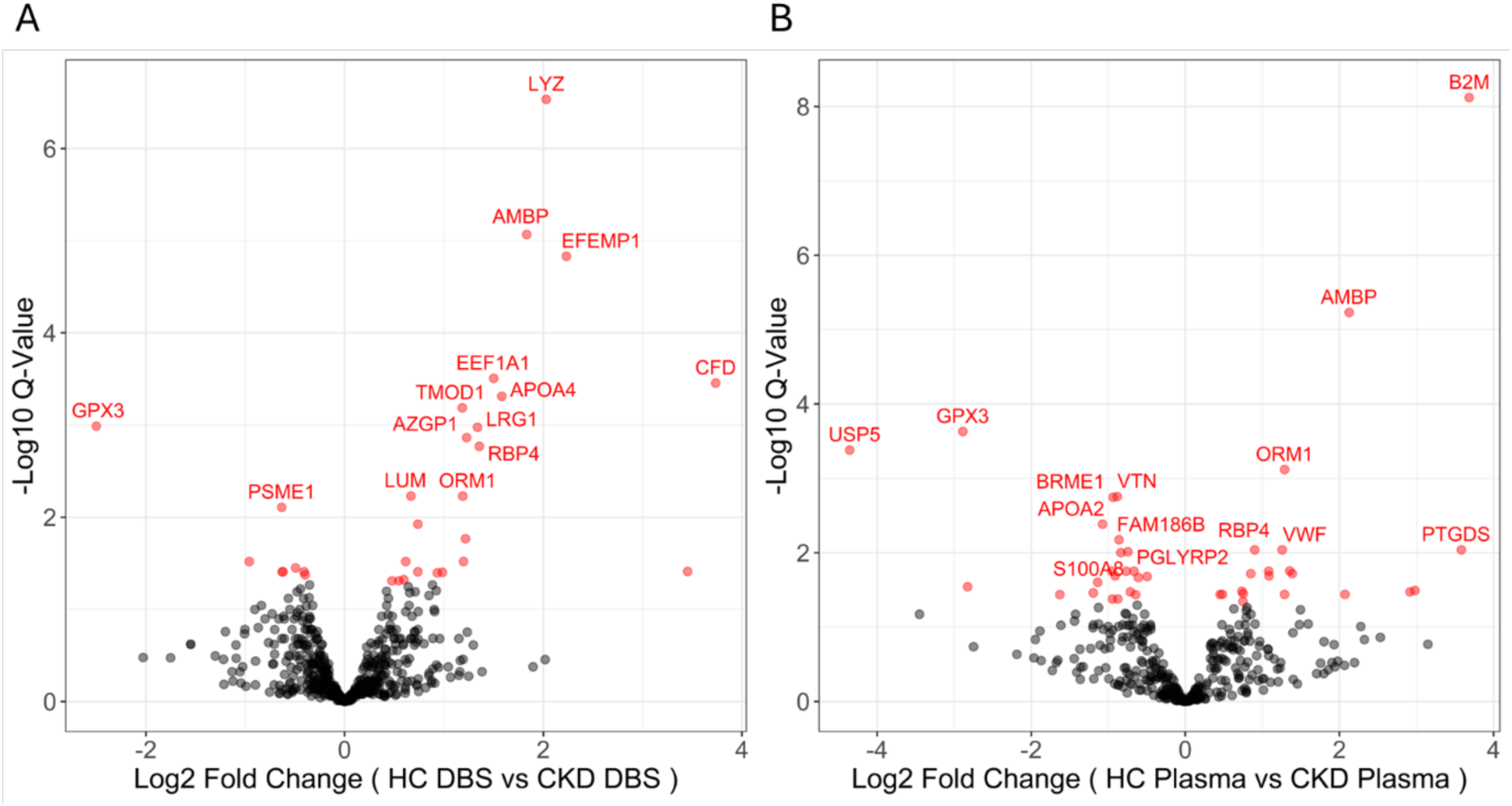
Quantitative profiling of dried blood spot proteomes. (A) Volcano plot displaying log2mean fold change in protein abundance against Q-values derived from 2-Sample significance tests (limma) comparing the identified proteomes from dried-blood spot (DBS) samples from healthy controls (HC) and CKD patients. Symbols corresponding to proteins with the greatest statistical confidence and highest fold change are labelled. (B) Volcano plot displaying log2mean fold change in protein abundance against Q-values derived from 2-Sample significance tests (limma) comparing the identified proteomes from plasma samples from healthy controls (HC) and CKD patients. Symbols corresponding to proteins with the greatest statistical confidence and highest fold change are labelled. (Protein names correlating to the Gene Name identifications can be found in the Supplementary information (supplementary table 2 and 3).

As expected, fewer proteins were identified from the CKD5 and healthy control plasma samples (474±16 and 477±14 respectively), compared with the DBS. Of these, 42 were significantly different between the two clinical cohorts (Figure 5B, Supplementary Sheet 10), with 20 (48%) being elevated in CKD5, although this did not include CRP, which was not significantly different between the two cohorts (Q-value= 4.2x10^-1^). Interestingly, both haemoglobin subunits HBG1 and HBG2 were increased in CKD5 patients (HBG1 log2FC= 2.98, Q-value= 3.2x10^-2^; HBG2 log2FC= 2.92, Q-value= 3.4x10^-2^), presumably due to the complex biological changes seen in relation to CKD associated anaemia, or as a consequence of the haemodialysis intervention on the blood.

Of the 73 proteins identified in total as significantly different between disease and control, 8 were identified in both the DBS and plasma samples (6 upregulated in CKD). Many of these have well-established roles in renal physiology and kidney disease. The upregulated proteins include: Alpha-1-microglobulin (AMBP) (DBS log2FC= 1.83, Q-value= 8.6x10^-6^; Plasma log2FC= 2.13, Q-value= 5.9x10^-6^), Retinol Binding Protein 4 (RBP4) (DBS log2FC= 1.36, Q-value= 1.7x10^-3^; Plasma log2FC= 0.90, Q-value= 9.1x10^-3^), Alpha-1-acid-glycoprotein (ORM1) (DBS log2FC= 1.19, Q-value= 5.9x10^-3^; Plasma log2FC= 1.29, Q-value= 7.6x10^-4^), Complement Factor D (CFD) (DBS log2FC= 3.74, Q-value= 3.5x10^-4^; Plasma log2FC= 2.07, Q-value= 3.6x10^-2^), Lysozyme C (LYZ) (DBS log2FC= 2.03, Q-value= 2.9x10^-7^; Plasma log2FC= 0.75, Q-value= 3.5x10^-2^), and Tropomodulin 1 (TMOD1) (DBS log2FC= 1.18, Q-value= 6.5x10^-4^; Plasma log2FC= 0.85, Q-value= 1.92x10^-2^).

AMBP is produced in the liver and adipose tissue and is known to degrade reactive oxygen species protecting tissues from damage. It is filtered in the glomeruli and degrades in the proximal tubular cells, where it can also be reabsorbed (64). Although the link between increased AMBP expression and CKD is not fully understood, it is theorised that increased oxidative stress in CKD upregulates AMBP, due to its antioxidative nature (65). It serves as a sensitive marker of tubular injury and oxidative stress in kidney disease (66). A similar hypothesis is used to explain ORM1 upregulation, which is an acute-phase protein, shown to increase in cases of inflammation, and known to protect against oxidative stress (67). It has been linked to renal injury, and is thought to correlate with disease severity (68). The precise role of RBP4, and how it may contribute to the pathophysiology of CKD are not clearly elucidated, however it is circulated by the kidneys and catabolised in the renal tubules, therefore reduction in kidney function is thought to lead to RBP4 accumulation and build up within the blood (69). Elevated levels of circulated or urinary RBP4 are therefore proposed to correlate with reduced glomerular filtration rate, tubular disfunction, and CKD progression (70). TMOD1 is an actin capping protein and assists in maintaining the structure of the distal tubules of the kidney, therefore helping to regulate water homeostasis. Disruption to healthy TMOD1 levels could therefore indicate issues with kidney function, due to its integral role (71). Additionally, LYZ and CFD are both involved in the innate immune response, activation of which is heavily associated with CKD pathogenesis and progression (72, 73, 74).

Alternatively, the two proteins shown to be downregulated in both the DBS and plasma samples are both known to protect against oxidative stress. ATP-citrate lyase (ACLY) (DBS log2FC= -0.40, Q-value= 4.22x10^-2^; Plasma log2FC= -0.91, Q-value= 2.0x10^-2^), and Glutathione peroxidase 3 (GPX3) (DBS log2FC= -2.50, Q-value= 1.0x10^-3^; Plasma log2FC= -2.89, Q-value= 2.3x10^-4^). ACLY is often seen to be downregulated in chronic diseases such as CKD due to metabolic alterations, such as the Warburg effect, leading to dyslipidaemia, and as a result lower levels of ACLY (75). The reduction in GPX3 levels, however, is likely due to imbalance between its production and usage rate. GPX3 is produced in the kidneys, a process which slows with kidney dysfunction (as is seen in CKD), leading to comparatively decreased levels. Additionally, GPX3 catalyses the reduction of reactive oxygen species, a process often referred to as scavenging. As a result, GPX3 can be transiently oxidised, meaning that at any given time point there may be variable levels of GPX3 identifiable (76).

Complement proteins were also observed to be elevated in CKD, irrespective of sampling strategy, however the specific complement proteins showing significance varied dependent on sampling type. The plasma samples showed significant elevation of C2 (Q-value= 3.6x10^-2^), C4BPB (Q-value= 1.8x10^-2^), C5 (Q-value= 3.6x10^-2^), CFD (Q-value= 3.6x10^-2^), and CFI (Q-value= 3.3x10^-2^), whilst DBS showed significant elevation of CFD (Q-value= 3.5x10^-4^), and CFHR2 (Q-value= 4.0x10^-2^). The complement system has established roles in immune mediated kidney diseases, such as lupus nephritis, atypical haemolytic uremic syndrome and C3 glomerulopathy, which can all lead to the development of CKD, however the role of complement in CKD5 is not clearly elucidated (77, 78).

Additionally, there were 23 proteins only identified as significantly different between the healthy control and CKD cohorts using the DBS samples, six of which have been shown to directly link to CKD and renal function (including CRP, which has been previously discussed): Fibulin 3 (EFEMP1) (log2FC= 2.23, Q-value= 1.5x10^-5^), Leucine-rich alpha-2-glycoprotein 1 (LRG1) (log2FC= 1.34, Q-value= 1.0x10^-3^), Lumican (LUM) (log2FC= 0.67, Q-value= 5.9x10^-3^), Defensin Alpha 1 (DEFA1) (log2FC= 0.60, Q-value= 4.8x10^-2^), and Apolipoprotein A-IV (APOA4) (log2FC= 1.58, Q-value= 4.9x10^-4^).

EFEMP1 is an extracellular matrix glycoprotein involved in tissue structural organisation and remodelling and is associated with declining kidney function. It is hypothesised to contribute to the progression of renal impairment (79). Similarly, LRG1 is a multifunctional signalling protein implicated in fibrosis and angiogenesis. It is consistently upregulated across various kidney diseases, where it contributes to pathological TGF-β signalling and progression of renal fibrosis (80). LUM represents another extracellular matrix-associated protein, which is known to be upregulated in diabetic nephropathy and to correlate inversely with estimated glomerular filtration rate (eGFR) (81). Additionally, DEFA1, is an antimicrobial protein involved in innate immune defence within the kidney. It contributes to host protection against infection, whilst also modulating inflammatory responses (82). Finally, APOA4 is an apolipoprotein involved in lipid metabolism, that has been identified as a circulating biomarker predictive of diabetic kidney disease progression. Its levels are associated with metabolic dysfunction and reported to reflect early signs of kidney disease progression (83).

Overall, the proof-of-principle cohort identified proteomes from DBS and plasma samples that were comparable, with all common statistically significant proteins exhibiting the same directionality of change, increasing their confidence as CKD markers. However, it is worth noting that there were differences in the relative abundance of these changes between the sample types.

## Conclusions

In this study, we establish and validate an optimised workflow for LC–MS/MS-based proteomic analysis of dried blood spots (DBS), with a particular focus on improving proteome depth through haemoglobin depletion and evaluation of a novel cell-free DBS device. Systematic evaluation of reconstitution buffers and DBS devices demonstrated that 50 mM ammonium bicarbonate provided optimal and reproducible protein recovery. Across the evaluated workflows, depletion using HemoVoid resulted in the greatest performance, increasing protein identifications from 364 ± 5 in undepleted samples to 545 ± 5, alongside improved reproducibility, with 94.7% of proteins observed across all replicates. This improvement is consistent with the reduction in haemoglobin content to 24 ± 7.9% of the undepleted level, supporting the importance of dynamic range reduction for enhanced proteome coverage.

Comparison of DBS with matched plasma and dried plasma samples confirmed that DBS provides substantially greater proteome coverage, with 573 ± 2 proteins identified compared to 181 ± 3 and 209 ± 3 for wet and dried plasma, respectively. This increase reflects the contribution of cellular proteins, supported by gene ontology analysis indicating enrichment of cytosolic components. Despite these compositional differences, drying itself introduced minimal bias, with no significant difference between wet and dried plasma proteomes.

Evaluation of cell-free DBS devices demonstrated that plasma-like samples can be obtained without post-collection depletion, being comparable in terms of proteome coverage to wet plasma, but lower with respect to conventional DBS given the absence of cellular material. These findings highlight a trade-off between analytical depth and workflow simplicity. Cell-free devices reduce processing time and cost, with an approximately four-fold cost saving compared to depletion-based approaches and are therefore well suited to large-scale studies where throughput and cost efficiency are prioritised. In contrast, conventional DBS combined with haemoglobin depletion provides substantially greater proteome coverage and is more appropriate for discovery-driven applications where detection of lower abundance or intracellular proteins is required.

Application of the optimised workflow to a cohort of patients with stage 5 CKD identified 31 differentially abundant proteins compared with matched controls. Clinically relevant markers such as C-reactive protein were significantly elevated in DBS from patients, demonstrating that DBS sampling captures known disease-associated biology. Plasma analysis yielded fewer protein identifications overall (∼30%) than DBS, with a partially overlapping set of significant proteins being identified demonstrating the same trends in differential abundance. These results demonstrate that DBS and plasma provide complementary views of disease biology, with DBS offering additional sensitivity to whole-blood derived changes. While we have demonstrated proof of principle here with data-dependent acquisition, proteome coverage will undoubtedly be improved in future investigations using data-independent acquisition workflows.

From a practical perspective, DBS sampling offers patients clear benefits as an adjust to traditionally collected samples for biobanking, including reduced invasiveness, preservation of venous access, and the ability to perform remote and longitudinal sampling. These advantages are particularly important in chronic conditions such as chronic kidney disease, where repeated sampling is required, or rare diseases which may require global participation. The choice of workflow should therefore be guided by both analytical and practical considerations. Depletion-based DBS workflows are advantageous when maximal proteome depth is required, for example in biomarker discovery or mechanistic studies. In contrast, cell-free DBS approaches are advantageous in population-scale or longitudinal studies where reduced cost, simplified processing, and patient convenience are prioritised. Moreover, the methods developed in this study are species independent and could therefore be used for a range of pre-clinical or veterinary research applications.

Collectively, this work demonstrates that DBS sampling, when combined with optimised processing, provides a robust and flexible platform for routine clinical proteomics. The ability to tailor workflows according to analytical depth, cost, and patient burden supports broader application across diverse study designs and clinical contexts and facilitates expanded access to large and geographically dispersed cohorts.

## Acknowledgements

A.J.C acknowledges funding from a Kidney Research UK Senior Non-Clinical Fellowship (SF_001_20221129) and funding from a Tenure Track Fellowship from the Institute of Systems, Molecular and Integrative Biology, Faculty of Health & Life Sciences, University of Liverpool. This research was also funded by Kidney Research UK (Paed_RP_008_20 190 926), Meditech Grant, and FAIR Ann Wyn Sherman Research Award (166185) (Funding Autoimmune Research Charity; registered UK Charity; number: 1176388) which was carried out at the National Institute for Health Research (NIHR), Alder Hey Children’s Research Facility. ED acknowledges clinical PhD funding from Wellcome Trust TRAP award and Alder Hey Children’s Kidney Charity. The views expressed are those of the author(s) and not those of the funders, NHS, the NIHR, or the Department of Health. This work was further supported by the UK’s Experimental Arthritis Treatment Centre for Children (supported by Versus Arthritis, Alder Hey Children’s NHS Foundation Trust, the Alder Hey Charity and the University of Liverpool). This research forms part of the LifeArc-KRUK Translational Centre for Rare Kidney Diseases (ref: 10749). The authors acknowledge use of The Liverpool University Biobank provided by the Faculty of Health and Life Sciences, University of Liverpool.

