## Supplementary Information for "Optimised haemoglobin depletion improves clinical proteomics from dried blood spots"

### Materials and Methods

Unless otherwise specified, all reagents were sourced from Thermo Fisher Scientific, Waltham, MA, United States.

#### *Haemoglobin Depletion*

HemogloBind Depletion was performed as per manufacturer's instructions, with 20 µL of reconstituted blood and 200 µL 0.02 M Dipotassium Phosphate (pH 6.5) added to a spin-filter tube, and vortexed for 5 minutes. The HemogloBind suspension (Biotech Support Group, South Brunswick, NJ, United States) was vortexed, and 200 µL added, and vortexed at RT for 10 minutes. The sample was centrifuged for 4 minutes at 15000 *g*.

NuGel-HemogloBind Depletion was performed as per manufacturer's instructions, with 50 mg of NuGel-HemogloBind Beads (Biotech Support Group, South Brunswick, NJ, United States) added to a spin-filter tube and conditioned with 200 µL of haemoglobin binding buffer (HBB) (Biotech Support Group, South Brunswick, NJ, United States). The tube was vortexed for 2 minutes, followed by centrifugation at 15000 *g* for 4 minutes. The filtrate was discarded. To a separate tube, 20 µL of blood was diluted in 400 µL of HBB and vortexed for 3 minutes, before being added to the beads. The mixture was vortexed for 10 minutes, followed by centrifugation at 15000 *g* for 4 minutes.

HemoVoid depletion was performed as per manufacturer's directions, with all measurements halved to account for reduced blood volume. Therefore, 25 mg of HemoVoid matrix (Biotech Support Group, South Brunswick, NJ, United States) was added to a microfuge tube, with 125 µL of haemoglobin binding buffer (HBB) (Biotech Support Group, South Brunswick, NJ, United States). The tube was vortexed, followed by centrifugation at 15000 *g* for 2 minutes. The filtrate was discarded, and 125 µL of HBB was added. The vortex and centrifuge steps were then repeated. An additional 150 µL of HBB, and 150 µL of blood sample was added. The sample was vortexed for 10 minutes, followed by centrifugation at 15000 *g* for 4 minutes. The filtrate was removed, and 250 µL of wash buffer (Biotech Support Group, South Brunswick, NJ, United States) was added. The sample was vortexed for 5 minutes, followed by centrifugation at 15000 *g* for 4 minutes, and the filtrate was discarded as wash. The washing step was repeated twice more. To the beads, 150 µL of elution buffer (Biotech Support Group, South Brunswick, NJ, United States) was added, and the sample was vortexed for 10 minutes. Finally, the sample was centrifuged for 4 minutes at 15000 *g*. In all cases the depleted sample was the final filtrate.

#### *Protein Digestion*

SP3 digestion was performed in a semi-skirted 96-well plate, using 25 µL of depleted blood sample. Samples were reduced and alkylated with 10 µL of Tris(2-carboxyethyl)phosphine (TCEP)/Chloracetamide (CAA) (120 mM/480 mM). After incubation for 5 minutes at 95 °C, the plate was centrifuged at 300 *g* for 1 minute then 10 µL of washed and combined Sera-Mag SP3 beads (E3 and E7) (Cytiva, Danaher, Washington DC, USA) was added. Next, 130 µL 100% ethanol were added and mixed. The plate was incubated for 10 minutes, before a magnetic rack was used to pellet the beads, and the supernatant was removed. The pellets were resuspended in 300 µL of 80% (v/v) ethanol and water, and, using a magnetic rack to pellet the beads, the supernatant was removed. This washing process was repeated for a total of six times. The pellets were dried using the SpeedVac (Centrifuge: UNIVAPO – 150 ECH, Cooling unit: UNICRYO MC2L -60 °C, Vacuum pump: UNIVAC DQ4) for 10 minutes. Samples were resuspended in 150 µL 100 mM Ammonium Bicarbonate with 300 ng LysC (Promega, Madison, WI, United States), and incubated in a ThermoMixer for 2 hours at 37 °C and 1450 rpm. Trypsin (Promega, Madison, WI, United States) was then added at 50:1 Protein:Trypsin, and the plate was incubated overnight at 37 °C and 1450 rpm. A magnetic rack was used to pellet the SP3 beads, and the supernatant was transferred to a new 96-well plate before being acidified with 5 µL 10% TFA and dried using the SpeedVac.

***LC-MS/MS***

EvoTips (EvoSep, Odense, Denmark) were prepared as per the manufacturer's instructions. They were washed with 20 µL 0.1% (v/v) FA (ACN) and centrifuged at 800 *g* for 1 minute. The tips were then conditioned using isopropyl alcohol, before washing with 20 µL 0.1% FA (v/v) ($H_2O$), and centrifugation at 800 *g* for 1 minute. Next 20 µL of sample (100 ng final peptide mass injected) was added, and the centrifugation step was repeated. The tips were then washed once more with 0.1% FA ($H_2O$), and centrifuged. To prevent the tips from drying out before analysis, 100 µL 0.1% FA ($H_2O$) was then added. Samples were ran with the following method https://www.evosep.com/wp-content/uploads/2021/05/AN-009A-30SPD.pdf

### Results and Discussion

#### *Optimisation of Sample Reconstitution Buffer*

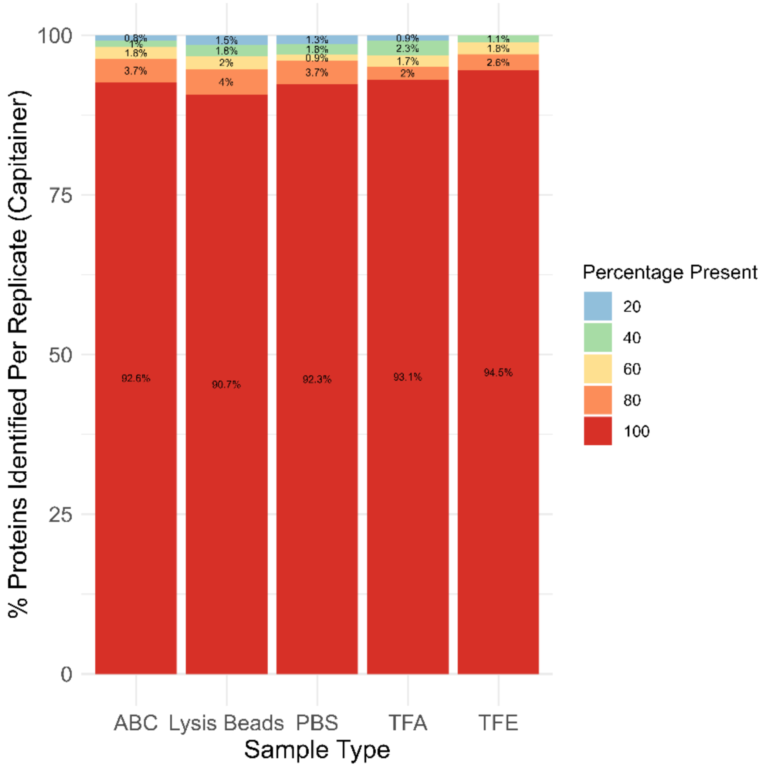

*Supplementary Figure 1- Optimisation of Sample Reconstitution Buffer for Capitainer b10 devices. Bar chart showing the reproducibility of protein identifications using each reconstitution buffer (n= 5) with the Capitainer b10 devices.*

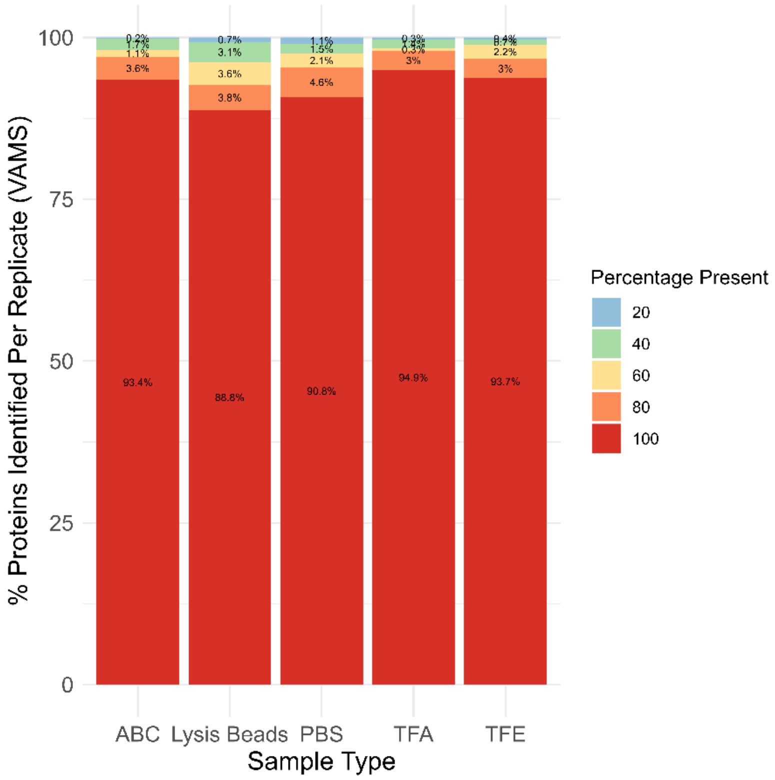

*Supplementary Figure 2- Optimisation of Sample Reconstitution Buffer for VAMS Mitra devices. Bar chart showing the reproducibility of protein identifications using each reconstitution buffer (n= 5) with the VAMS Mitra devices.*

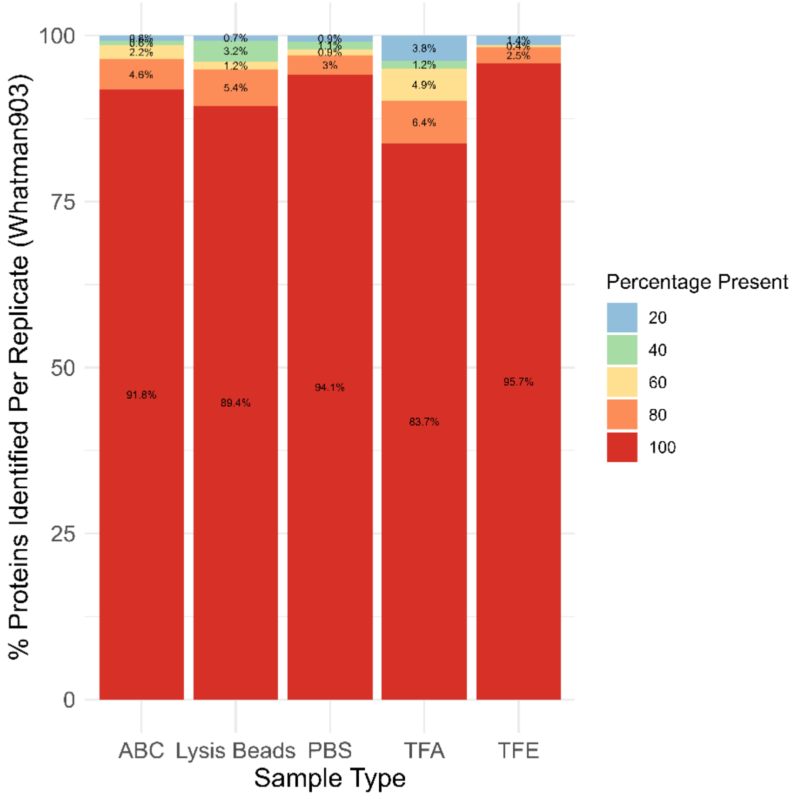

*Supplementary Figure 3- Optimisation of Sample Reconstitution Buffer for Whatman903 devices. Bar chart showing the reproducibility of protein identifications using each reconstitution buffer (n= 5) with the Whatman903 devices.*

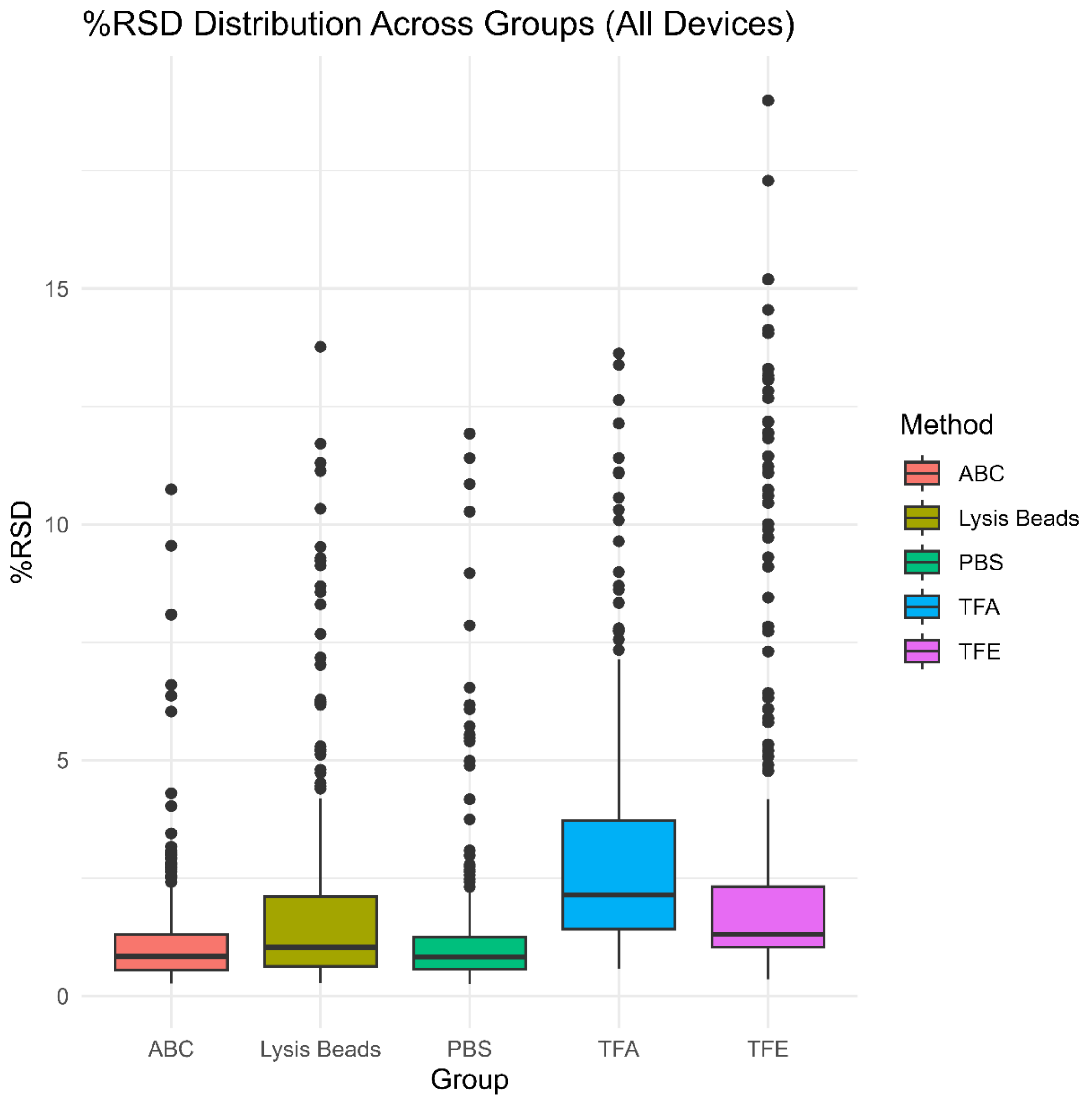

*Supplementary Figure 4- Optimisation of Sample Reconstitution Buffer. Bar chart showing the percentage relative standard deviation of protein relative abundance using each reconstitution buffer across all 3 devices (n=15).*

***Evaluation of High Abundance Protein Depletion Techniques***

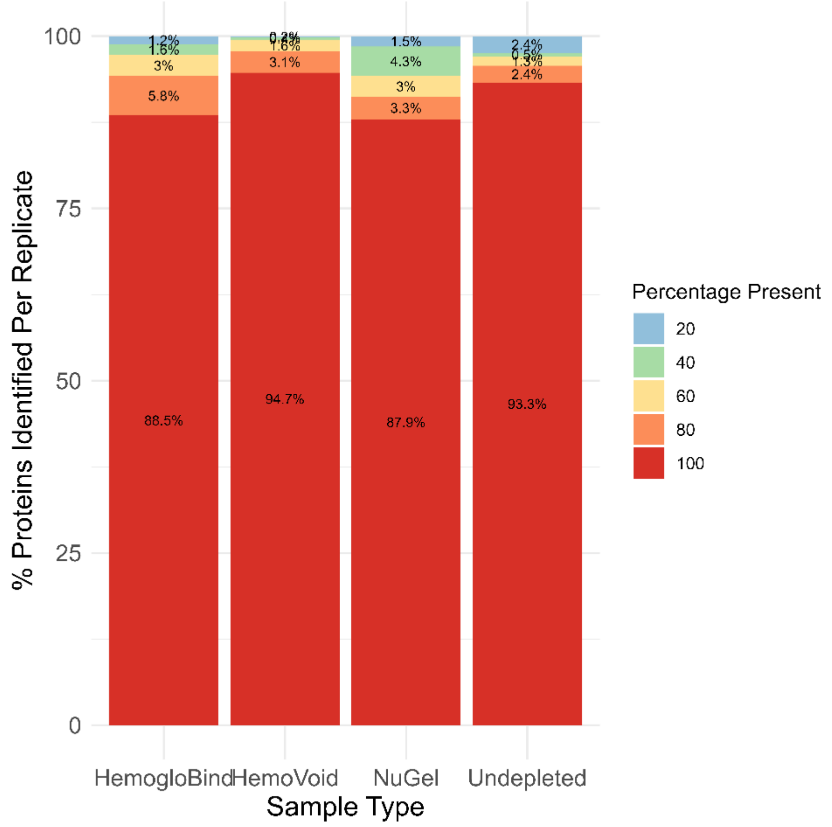

*Supplementary Figure 5- Evaluation of High Abundance Protein Depletion Techniques. Bar chart showing the reproducibility of protein identifications using each depletion technique (n= 5).*

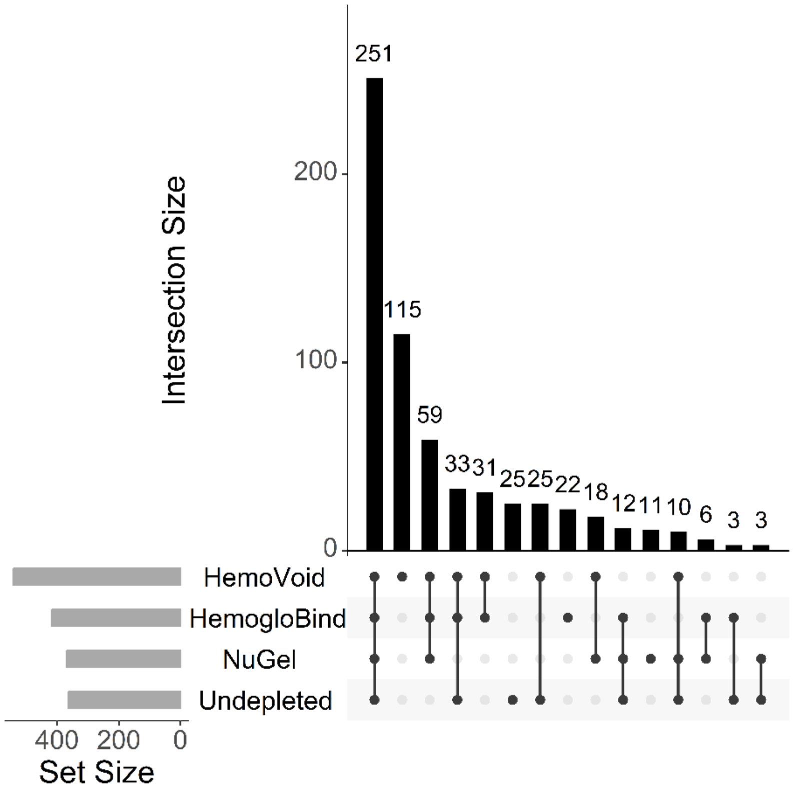

*Supplementary Figure 6- Evaluation of High Abundance Protein Depletion Techniques. Upset plot showing the cross over of proteins identified using each depletion method (n= 5)*

***Cell Free Dried Blood Spot Analysis***

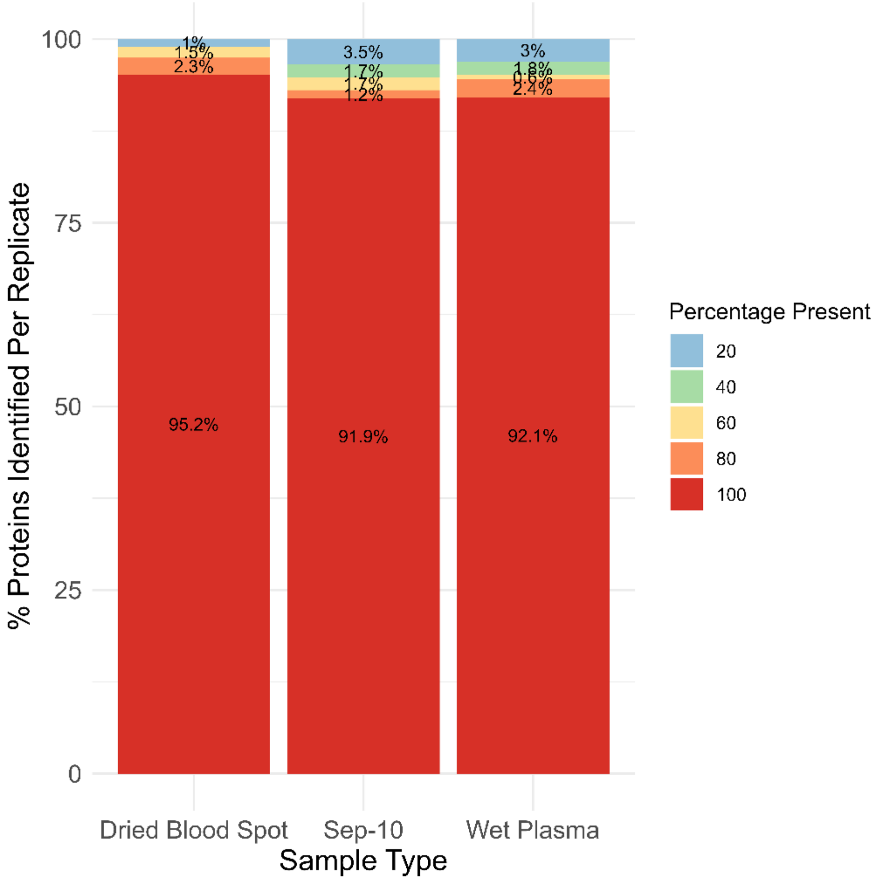

*Supplementary Figure 7- Cell Free Dried Blood Spot Analysis. Bar chart showing the reproducibility of protein identifications using each sampling method (n= 5).*
